# Are phylogenetic patterns the same in anthropology and biology?

**DOI:** 10.1101/006486

**Authors:** David A. Morrison

## Abstract

The use of phylogenetic methods in anthropological fields such as archaeology, linguistics and stemmatology (involving what are often called “culture data”) is based on an analogy between human cultural evolution and biological evolution. We need to understand this analogy thoroughly, including how well anthropology data fit the model of a phylogenetic tree, as used in biology. I provide a direct comparison of anthropology datasets with both phenotype and genotype datasets from biology. The anthropology datasets fit the tree model approximately as well as do the genotype data, which is detectably worse than the fit of the phenotype data. This is true for datasets with <500 parsimony-informative characters, as well as for larger datasets. This implies that cross-cultural (horizontal) processes have been important in the evolution of cultural artifacts, as well as branching historical (vertical) processes, and thus a phylogenetic network will be a more appropriate model than a phylogenetic tree.

## 1. Introduction

There has been much talk over the past few decades about the extent to which the various disciplines within anthropology (in the broadest sense) can use, or benefit from, methodological techniques developed in other disciplines, notably biology (see Mace et al. 2005; Forster & Renfrew 2006; Lipo et al. 2006). This has been particularly true for historical studies of languages (linguistics) and texts (stemmatology), past cultures (archaeology), and physical type (physical */* biological anthropology). This is a debate to which both biologists and anthropologists can contribute. Here, I investigate the extent to which anthropological and biological datasets fit the same mathematical model, and therefore the extent to which traditional phylogenetic methods might apply to anthropology.

As noted by Nunn (2011), evolutionary anthropology involves studies such as assessing the taxonomic affinities of newly discovered hominins, reconstructing primate phylogeny, constructing language trees, evaluating the history of copying of written texts or oral tales, and the evolution of tools and other manufactured items. All of these involve the techniques associated with the comparative method, of which phylogenetics is one. We therefore need to quantitatively evaluate the extent to which phylogenetic techniques, as developed in biology, can be applied to anthropology.

The use of phylogenetic methods seems to be relatively unproblematic in the case of physical anthropology (i.e. studies of the origin and evolutionary development of humans as a species; Holliday 2003), although this particular field is concerned as much with population genetics as it is with species phylogenies (especially the emerging field of population genomics). However, the use of phylogenetic methods in other anthropological fields, such as archaeology, linguistics and stemmatology (involving what are often called “culture data”), is based on an analogy between human cultural evolution and biological evolution (Mace & Holden 2005; Pagel 2009; Howe & Windram 2011).

This analogy assumes that the patterns of historical change in anthropology and biology are similar enough that the analytical methods can be combined. It is not actually a requirement, but there is often the assumption that the underlying processes are also similar. Both anthropology and biology apparently involve an evolutionary process, in which the study objects form groups that change via modification of their intrinsic attributes, the attributes being transformed through time from ancestral to derived states (often called “innovations” in anthropology). However, this apparent similarity is basically a metaphor, because human culture is not a collection of biological objects. In Popperian terms, biology is part of the “world that consists of physical bodies” while culture and linguistics are part of the “world of the products of the human mind” (Geisler & List 2013).

If we are going to draw an analogy between anthropological studies and biological studies, and use this analogy to justify the use of certain analytical techniques, then we need to understand the analogy thoroughly. One important aspect of this analogy is how well anthropological data fit the model of a phylogenetic tree, as used in biology, and this is what I investigate here.

## 2. The tree model in anthropology

It has often been claimed that cross-cultural processes, such as trade and exchange, have been more important in the evolution of cultural ideas and artifacts (in the broadest sense) than have branching historical processes. That is, horizontal processes dominate rather than vertical processes, as has been traditionally assumed in phylogenetics (especially for eukaryotes). Under these circumstances, the data would fit a network model rather than a tree model, the latter being the basis of most phylogenetic analyses (Felsenstein 2004). If so, then the tree-based phylogenetic methods would be of limited practical use in anthropology.

As a quantitative contribution to the debate, Collard et al. (2006) compared 21 anthropological datasets with 21 biological datasets, to assess the extent to which they fit a phylogenetic tree model. They analyzed each dataset using maximum parsimony, and then calculated the ensemble retention index (RI), which ranges from 0 (none of the characters are compatible with the optimal tree) to 1 (all of the characters are compatible with that tree). Their conclusion was that the range of index values did not differ between the anthropological and biological datasets, and so the anthropological data fit a tree model at least as well as do the biological data.

Unfortunately, there are two problems with the datasets used by Collard et al., and these biases combine to invalidate their conclusion. First of all, among their datasets there is a clear relationship between the RI and the number of parsimony-informative characters, and the datasets with the greatest number of such characters were all biological. Second, the biological datasets included both phenotypic and genotypic characters, and there is no a priori reason to believe that the genotypic datasets are directly comparable to the anthropological ones.

These two problems are illustrated in Figure 1. Among the 42 datasets, the log-linear correlation between the RI and the number of parsimony-informative characters is r = −0.502 (p = 0.019), indicating that 25% of the variation in the RI is accounted for solely by variation in the number of parsimony-informative characters. Unfortunately, all of the datasets with >200 parsimony-informative characters were biological; and this imbalance has a marked influence on the comparison between the two sources of data. (Note that in counting the characters I have made no attempt to distinguish between datasets with multi-state characters and those where the data have been coded as binary. This could be done by re-coding all of the multi-state characters as binary or additive binary. Doing so would refine the analysis but not change the conclusions.)

**Fig. 1.**
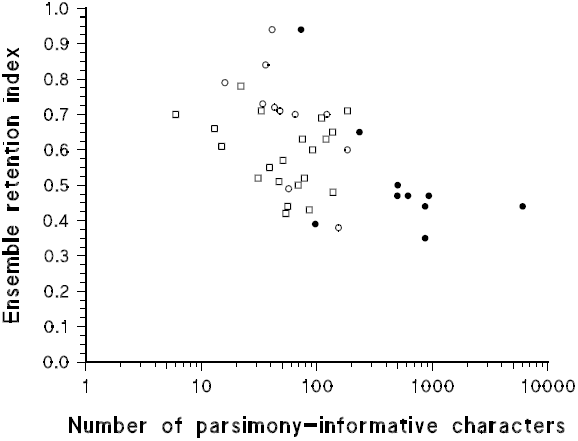
Scatterplot of the relationship between the (ensemble) retention index and the number of parsimony-informative characters for 21 empirical datasets from biology and 21 datasets from anthropology. Open circles: phenotype data; closed circles: genotype data; open squares: culture data. The data are taken from Collard et al. (2006).

Furthermore, 11 of the biological datasets consisted of phenotype data alone (morphology or behavior) and 10 of them consisted of genotype data alone (mtDNA sequences). The graph (Fig. 1) shows that the genotype datasets have, in general, more parsimony-informative characters and a smaller RI than do the phenotype datasets. This makes inappropriate any comparison with the anthropological data (consisting of human behavior, and the morphology of cultural artifacts), due to bias. A Welch t-test (which accounts for the unequal variances of the two samples) comparing the RIs of the anthropological datasets to all of the biological datasets yields t = 0.426 (p = 0.673). This inappropriate test apparently supports the conclusion of Collard et al., that the anthropological datasets fit a tree model just as well as do the biological datasets. However, a Welch t-test more appropriately comparing the anthropological datasets solely to the phenotype biological datasets yields t = 2.011 (p = 0.063), which suggests that the anthropological datasets might fit a tree model less well than do the biological datasets.

Clearly, we need a revised comparison in order to validly test the original hypothesis proposed by Collard et al.

## 3. Revised comparison

My revised comparison consists principally of expanding the collection of phenotype biological datasets, in order to make a valid comparison of them to the anthropological datasets. However, I also increase the total sample size somewhat, and I also make a direct comparison between the genotype biological datasets and some of the anthropological datasets.

The extra biological phenotype datasets were obtained by searching the TreeBASE 2.0 database (Sanderson, et al. 1994) for studies with a title containing the words “morphology” or “behavior” / “behaviour”. All 20 studies retrieved for the Vertebrata and Insecta that actually contained some non-genotype data were used (Table 1). (Note that the taxonomic restriction follows the choice of datasets of Collard et al.) All of these datasets have <500 parsimony-informative characters. Five genotype datasets with >1000 parsimony-informative characters were also used, taken from previous comparative studies that I have conducted (Table 1).

**Table 1.** Datasets analyzed, in addition to those listed by Collard et al. (2006)

| Dataset | Number of taxa | Number of characters | Parsimony-informative characters | Retention index | Source <sup>a</sup> |
| --- | --- | --- | --- | --- | --- |
| **Biology — phenotype** |  |  |  |  |  |
| Mammal anatomy | 63 | 439 | 417 | 0.76 | Phillips et al. (2009) |
| Cetacea anatomy | 28 | 9 | 9 | 0.81 | Thewissen & Madar (1999) |
| Crocodylia anatomy | 62 | 164 | 158 | 0.82 | Brochu (1997) |
| Iguanid anatomy | 13 | 89 | 68 | 0.84 | Vieira et al. (2005) |
| Ornithischian dinosaur anatomy | 14 | 221 | 124 | 0.58 | Norman et al. (2011) |
| Ornithischian dinosaur anatomy | 12 | 10 | 10 | 0.91 | Sereno (2010) |
| Amphibia anatomy | 49 | 161 | 159 | 0.71 | Laurin & Vallin (2004) |
| Amphibia anatomy | 16 | 24 | 23 | 0.68 | Grandison (1981) |
| Insect morphology | 26 | 176 | 127 | 0.83 | Whiting et al. (1997) |
| Hemiptera morphology | 66 | 83 | 68 | 0.83 | Cryan et al. (2004) |
| Hymenoptera morphology | 44 | 57 | 54 | 0.65 | Weiblen (2001) |
| Hymenoptera morphology | 11 | 15 | 11 | 0.90 | Lee et al. (1992) |
| Hymenoptera morphology <sup>b</sup> | 65 | 109 | 109 | 0.69 | Alexander & Michener (1995) |
| Hymenoptera morphology | 73 | 88 | 78 | 0.86 | Heraty et al. (2004) |
| Lepidoptera morphology | 47 | 236 | 208 | 0.73 | Nazari et al. (2007) |
| Neuroptera morphology | 23 | 100 | 85 | 0.76 | Beutel et al. (2010) |
| Phthiraptera morphology | 16 | 55 | 46 | 0.74 | Banks & Peterson (2004) |
| Phthiraptera morphology | 122 | 58 | 58 | 0.81 | Page et al. (1995) |
| Phthiraptera morphology | 61 | 138 | 131 | 0.65 | Smith (2001) |
| Spider behavior | 22 | 12 | 12 | 0.72 | Hormiga et al. (1995) |
| **Biology — genotype** |  |  |  |  |  |
| Mammalia — 22 genes | 44 | 17,028 | 8,004 | 0.58 | Morrison (2007) |
| Mammalia — 18 genes | 66 | 10,694 | 4,924 | 0.43 | Morrison (2007) |
| Fungi — 5 genes | 106 | 3,317 | 1,852 | 0.31 | Morrison (2007) |
| Apicomplexa — 3 genes | 35 | 3,793 | 1,293 | 0.57 | Morrison (2006) |
| HIV-1 genome | 94 | 10,922 | 6,088 | 0.40 | Morrison (2007) |
| **Culture** |  |  |  |  |  |
| Behavior | 43 | 11 | 11 | 0.69 | Walker et al. (2012) |
| Thai Buddha statues | 42 | 17 | 17 | 0.69 | Marwick (2012) |
| Musical instruments | 39 | 17 | 15 | 0.80 | Tëmkin & Eldredge (2007) |
| Musical instruments | 49 | 59 | 48 | 0.54 | Tëmkin & Eldredge (2007) |
| Japanese languages | 19 | 350 | 185 | 0.75 | Lee & Hasegawa (2013) |
| Japanese languages | 59 | 675 | 366 | 0.60 | Lee & Hasegawa (2011) |
| Greek text | 166 | 761 | 435 | 0.61 | Spencer et al. (2002) |
| Finnish text | 52 | 1,185 | 296 | 0.50 | Roos & Heikkilä (2009) |
| Biblical text | 26 | 50 | 40 | 0.67 | Unpublished |
| Folk tales | 58 | 72 | 71 | 0.73 | Tehrani (2013) |
| **Linguistics and stemmatology — large datasets** |  |  |  |  |  |
| World languages | 101 | 2,301 | 982 | 0.75 | Dun et al. (2011) |
| Austronesian languages | 77 | 5,185 | 4,554 | 0.26. | Gray & Jordan (2000) |
| Indo-European languages | 103 | 6,280 | 2,560 | 0.70 | Bouckaert et al. (2012) |
| Norse text<sup>b</sup> | 43 | 3,318 | 2,063 | 0.63 | Robinson & O'Hara (1996) |

<sup>a</sup> Sources are listed in online Appendix 1.
<sup>b</sup> Information taken from the original publication.

Extra anthropological datasets (Table 1) were also taken from the literature (plus an unpublished one of my own). Four of these datasets match the types of study chosen by Collard et al., which were mainly from archaeology and physical anthropology. However, four studies were also specifically chosen from stemmatology, along with five studies from linguistics.

For each dataset, the search for the maximum-parsimony tree was conducted using the parsimony ratchet (Nixon 1999) with 200 replicates. These searches, and all statistical calculations, were conducted using PAUP* 4.0b 10 (Swofford 1998), and the ratchet commands were produced with PAUPrat (Sikes & Fewis 2001). The data are available in the Appendices (Supplementary Online Material).

The new data for the main comparison are illustrated in Figure 2a, with 31 datasets for both the phenotype biological data and the anthropological data, containing <500 parsimony-informative characters. For these data there is no correlation between the character number and the RI (r = - 0.164, p = 0.203). More to the point, there is a large difference in RI between the anthropological and biological datasets (Welch t-test: t = 4.502, p < 0.0001), with the anthropological data fitting a tree model much less well than do the biological data.

**Fig. 2.**
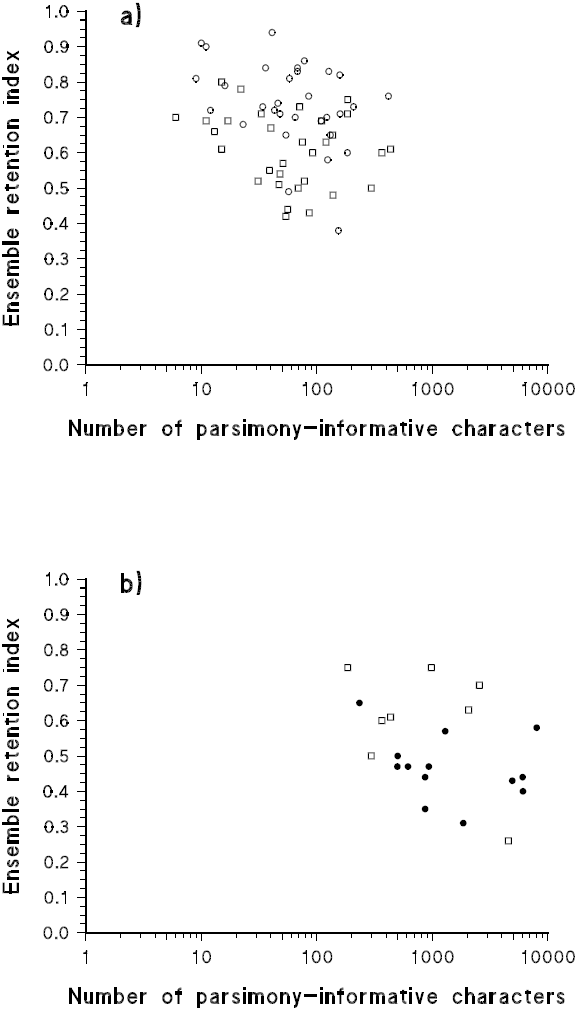
Scatterplot of the relationship between the ensemble retention index and the number of parsimony-informative characters for (a) 31 phenotype datasets from biology and 31 datasets from anthropology, and (b) 13 molecular datasets from biology and 8 datasets from linguistics and stemmatology. Open circles: phenotype data; closed circles: genotype data; open squares: culture data.

This leads to the issue of whether genotype biological data can validly be compared to anthropology data. I discuss the theory in the next section, but one practical problem is that genotype data often have many more characters than do anthropology data. There are, however, two anthropological fields in which thousands of characters can be collected: comparative linguistics and stemmatology. It is therefore at least possible to make a direct comparison with genotype data. I will briefly explain why these fields can produce such large datasets.

First, comparative linguistics is the study of the relationships between languages. Typically, lists of word meanings (a few hundred meanings) are recorded for the various languages being studied, and these lists are turned into cognate sets of putatively homologous words (a few thousand sets), which are then coded as a binary matrix (i.e. the multi-state data are re-coded as binary). This can lead to very large datasets; for example, Gray et al. (2009) produced a dataset with 400 languages and >34,000 cognate sets. Second, stemmatology is the study of the relationships between different versions of the same written text. The putatively homologous parts of the study texts are aligned and encoded as a string of characters, usually with each character corresponding to a word. The length of the alignment depends on the length of the text (hundreds to thousands of words). This can also lead to very large datasets; for example, Windram et al. (2008) produced a dataset with 21 texts and 16,548 characters.

Here, I have used 13 genotype datasets, from both the mitochondrial and nuclear genomes, including both protein-coding and RNA-coding sequences, and ranging from 234 to 8,004 parsimony-informative characters (Fig. 2b). For comparison, I have five datasets from linguistics and three from stemmatology, ranging from 185 to 4,554 parsimony-informative characters (Fig. 2b). There is a borderline statistical difference in the RIs of these two groups (Welch t-test: t = 2.096, p = 0.063), with the genotype datasets fitting the tree model less well than do the anthropology datasets.

So, while these are only small comparisons, there are two different patterns in the data. That is, in biology there is some indication that the large genotype datasets fit a tree less well than do the smaller phenotype datasets (mean RI ≈ 0.5 versus 0.7, respectively), whereas there is no difference in fit for the large versus small anthropological datasets (mean RI ≈ 0.6 in both cases). This possible distinction between genotype and phenotype data in biology might be worth investigating further.

## 4. Discussion

The answer to the question posed in the title of this paper is: apparently not, at least for datasets with <500 parsimony-informative characters. Under these circumstances, anthropological datasets fit the maximum-parsimony tree model less well than do phenotype biological datasets. This implies that, indeed, cross-cultural (horizontal) processes have been important in the evolution of cultural artifacts, as well as branching historical (vertical) processes, which is a different conclusion from that reached by Collard et al. This does not invalidate the use of phylogenetic analysis in culture studies, but it does mean that a phylogenetic network would be a more useful model than a phylogenetic tree (Bapteste et al. 2013).

Evolutionary theory has four basic contributions that it could possibly make to anthropology (Pappas & Mooers 2011): (i) correlating human genetics with other aspects of culture; (ii) adopting evolutionary theory to provide causal mechanisms in culture history; (iii) using evolutionary concepts as metaphors; and (iv) applying phylogenetic techniques to anthropological data. Here, we are interested in the use of contribution (iii) as a justification for adopting (iv). Analyzing cultural data using phylogenetics is thus often seen as predicated on the adoption of the tree metaphor for representing phylogenetic history.

The tree metaphor has a long history in cultural studies. For example, the first empirical use of this metaphor in a chronological context appears to have been that of Buffon (1755) in biology, Gallet (1800) in linguistics, and Collin & Schlyter (1827) in stemmatology, all of which long pre-date empirical use within the context of modern evolutionary theory (Čelakovský 1853 in linguistics; Mivart 1865 in biology). The tree is thus apparently a powerful as well as a useful metaphor, although its adoption in culture studies has been only sporadic (e.g. see Geisler & List 2013 for linguistics). It is for this reason that the tests carried out by Collard et al. (2006) and myself are of interest.

There are, however, clear limitations to the comparison undertaken here. Notably, the phenotype data were restricted to those from the Vertebrata and Insecta, with one exception (spider behavior). This was apparently adopted by Collard et al. based on the idea of comparing humans with related animals. However, it seems very likely that if the comparison were to be made using other systematic groups then the result would be different. The evolution of plants, for example, is usually considered to have involved extensive hybridization (Mallet 2005), so that plant datasets are less likely to fit a tree model than are animals. A similar argument applies to prokaryotes, which have long been subject to horizontal gene transfer. However, this does not negate the overall conclusion, which is that a tree model is unlikely to be particularly suitable for anthropology studies.

Furthermore, the comparison made here is rather simplistic, and deliberately so. Culture evolution is complex, and a tree is a very simple model. However, the hypothesis tested here is comparative, comparing the use of the tree model in biology and anthropology — I am evaluating the relative applicability of the model, not its realism. So, the test here asks solely how well the data fit a tree within a parsimony context. This has been the most common context in anthropology, to date, and so it was natural for Collard et al. to use a maximum-parsimony analysis, and to assess the fit of the data to the optimal tree. Using the ensemble retention index (Farris 1989) keeps the focus on the characters themselves, rather than on any aspect of the tree building. For simplicity, I have followed suit.

However, in practice, any statistic that quantifies the fit of data to a tree could be used, and any tree-building algorithm could be used. An alternative, but still simple, approach would be to make the comparison using a distance-based analysis (e.g. Bryant et al. 2006). For example, the fit of the data to a neighbor-joining tree could be compared to its fit to a neighbor-net network — this would provide a heuristic assessment of how much better the data fit a reticulating network compared to a tree.

A more sophisticated approach would involve the direct comparison of different phylogenetic models in a likelihood context. The limitation here is the current dearth of explicit alternative models in anthropology. This topic is being actively investigated in the context of phylogenetic networks, which add reticulations in the basic tree model (e.g. Yu et al. 2011), but explicit network models have rarely been developed in anthropology (see Dunn 2014). For example, Ross et al. (2013) note that in genetics there are specific measures of genetic distance but that in their anthropology study they can use only a generic distance measure (Jaccard in their case). A more general point of interest here concerns the applicability of phylogenetic analysis to anthropological studies. It seems to be commonly accepted by proponents that an analogy can be drawn between anthropological data and specifically genotype data in biology (Mace & Holden 2005; Pagel 2009; Howe & Windram 2011), and that this can be used to justify the application of phylogenetics. This seems to be based on a misconception.

In culture studies information is usually passed between “generations” via repeated teaching, learning or imitation, whereas genotype information is passed as DNA-coded information in genes. Languages must be learned anew by each user, as must tool making and artistic procedures, whereas each possessor of a gene inherits a physical copy of the original. Furthermore, the DNA “alphabet” is a very different thing from a language alphabet, for example.

The recording of cultural information in permanent forms, such as written works, does not invalidate this point. Rather, if emphasizes further differences between cultural history and genotype history — that cultural information can skip many generations, and also that it can be transmitted diagonally as well as vertically and horizontally (i.e. simultaneously from one culture to another as well as from the distant past to the present). Genes do not skip generations, at least not in nature.

So, genes are physical objects whereas memes (cultural replicators) are theoretical concepts. It is not necessarily obvious what the replicating units of language, for example, really are — equating phonemes with nucleotides seems rather superficial. Concepts such as phonemes seem much more closely related to concepts like phenotype than to genotype. Genotype refers to measurement of features relating solely to the genes, which are the units of inheritance. Phenotype, on the other hand, refers to features that are the product of genes + interactions between genes + interactions between genes and the environment. This is much closer to the concept of characters in cultural studies.

For genetic data the units of comparison are to some extent self-evident (e.g. DNA residues). However, phenotypic data have traditionally required careful study of qualitative features, in order to establish the basis for comparing like with like, including comparative anatomy, developmental studies, and so on. A similar situation pertains in anthropology, where elucidating the nature of the characters to be compared has traditionally occupied much of the time of anthropologists. As far as phylogenetic methods are concerned, for both phenotype and anthropology data establishing the character transformation series is more than half the battle (and its description occupies a large proportion of any publication), and the algorithmic tree-building is to some extent then simply a graphical representation of the congruent patterns among those transformation series (based on an objective and repeatable method).

One can, nevertheless, question whether all forms of anthropology data have a phenotype in the same conceptual way. For example, some anthropology data refer to physical objects (pottery, rugs, arrowheads, etc) while others refer to immaterial concepts, such as phonemes in linguistics. Immaterial concepts are much more difficult to compare in any meaningful way, because there is no actual object to study. For example, the characters that have been used for phylogeny reconstruction in linguistics include phonological, lexical and syntactic traits. None of these approximates to a genetic character in any way, although they are all features that are “observable” in an objective and repeatable manner. Furthermore, linguistic borrowing cannot be simply equated with introgression or horizontal gene transfer, because the physical processes are quite different.

So, there seems to be no a priori reason to expect anthropological data to be analogous to genotype data. However, no such theoretical objection appears necessarily to apply to an analogy between culture data and phenotype data. The patterns produced by cultural processes and phenotypic processes have much more in common than either does with genotypic processes.

In this sense, the discovery of a practical difference between the phylogenetic patterns of biology and anthropology is of importance. Tree-based phylogenetic analyses will apparently be of considerably less practical usefulness in anthropology than they have been in animal biology. At best, they might function as a heuristic tool, but even that is not certain. A phylogenetic tree is a graph, and graphs are excellent tools for exploratory data analysis. However, a phylogenetic tree is a very specific type of graph, with a precise interpretation in terms of representing evolutionary history. If the model underlying that interpretation is inappropriate, then the analysis is unlikely to be much use even as a heuristic.

It would be more productive to employ analysis techniques that accommodate both horizontal and vertical evolutionary processes. Indeed, the first empirical phylogenetic models in stemmatology (Collin & Schlyter 1827), linguistics (Gallet 1800) and biology (Buffon 1755) all actually display reticulation, rather than a bifurcating tree, and so this suggestion is a long-standing one.

## Acknowledgments

Thanks to Mattias List and Jamie Tehrani for helpful correspondence.

## Appendices

Appendix 1. List of data sources (spreadsheet)

Appendix 2. Nexus-formatted data files (text file)

Available at (911 KB): http://www.rjr-productions.org/Supplementary.html

